# Community adaptation to temperature explains abrupt soil bacterial community shift along a geothermal gradient on Iceland

**DOI:** 10.1101/2022.04.04.486966

**Authors:** James T. Weedon, Erland Bååth, Ruud Rijkers, Stephanie Reischke, Bjarni D. Sigurdsson, Edda Oddsdottir, Jurgen van Hal, Rien Aerts, Ivan A. Janssens, Peter M. van Bodegom

## Abstract

Understanding how and why soil microbial communities respond to temperature changes is important for understanding the drivers of microbial distribution and abundance. A unique area in Iceland, where soil temperatures have increased due to geothermic activity four years prior to sampling, creating a stable gradient of ambient to +40°C, allowed us to investigate the shape of the response to warming of soil bacterial communities, and their associated community temperature adaptation. We used 16S rRNA amplicon sequencing to profile bacterial communities, and bacterial growth based assays (^3^H Leu-incorporation) to characterize community adaptation using a temperature sensitivity index (SI, log (growth at 40°C/4°C)). Samples were taken from ≥9 levels of warming (covering almost up to +40°C above ambient soil temperature), under both grassland (*Agrostis capillaris*) and forest (*Picea sitchensis*) vegetation. The soils had very different community composition, but temperature adaptation was the same. Both diversity and community composition as well SI showed similar threshold dynamics along the soil temperature gradient. There were no significant changes up to soil warming of approx. 6-9 °C, beyond which all indices shifted in parallel. The consistency of these responses gives strong support for a decisive role for direct temperature effects in driving bacterial community shifts along soil temperature gradients.

## Introduction

Soil microbial communities and their response to changing temperatures have been the focus of extensive research in the past decade (Zhou et al., 2016; Oliverio et al., 2017). Community responses to temperature and other climate-related variables is a fundamental ecological pattern which must be understood to fully explain the drivers of microbial distribution and abundance (Zhou et al., 2016; Delgado-Baquerizo et al., 2018). Moreover, it has been proposed that including microbial dynamics in ecosystem models will improve our ability to predict responses of biogeochemical cycles to changing climate conditions (Todd-Brown et al., 2012; Wieder et al., 2013, 2015), even though the putative link between community composition *per se* and ecological function remains elusive in most soil environments (Prosser, 2012; Bier et al., 2015)

For both fundamental ecological understanding, and for applications to ecosystem modelling, sub-arctic and arctic environments are considered particularly important for the study of the relation between soil microbes and temperature (Wieder et al., 2015, 2019). Current and future climate warming is at a higher magnitude than other parts of the globe (Post et al., 2019), and temperature sensitivity of soil microbial activity is higher at low temperatures (Kirschbaum, 1995, 2000). There is thus a large potential for a positive climate feedback via warming-induced increases in net C release, which may be mediated by the activity and temperature sensitivity of soil microbial communities (Cavicchioli et al., 2019).

A range of techniques have been used to study the relationship between soil microbial communities and temperature, including laboratory incubations (Oliverio et al., 2017), sampling along climatic gradients (Yergeau et al., 2007), and in-situ climate manipulation experiments (Weedon et al., 2017). Although providing relevant insights, each of these approaches have a number of limitations. The large spatial scales necessary for gradient studies may introduce additional confounding factors related to soil and vegetation. In experimental warming studies practical considerations compel investigators to choose a small number of (usually small) temperature steps (De Boeck et al., 2015). Studying temperature responses of soil communities along geothermal gradients at local-scales (10 – 100s of metres) overcomes these limitations (O’Gorman et al., 2014). Potential biogeographic and large-scale edaphic confounders are held constant; and a range of temperatures can be studied allowing for the study of dynamics not possible with step-wise experimental set-ups.

A near-surface geothermal system that arose after an earthquake in SW Iceland provides a valuable model system for studying soil temperature effects on ecosystem processes (Sigurdsson et al., 2016). This system has been used to study the effects of warming on a wide range of processes in soil ecology and biogeochemistry (Walker et al., 2018, 2020; Marañón-Jiménez et al., 2019; Poeplau et al., 2020; Zhang et al., 2020) Focussing on the microbial community composition, Radujković et al. (2018) found that bacterial and fungal communities changed only at warming levels exceeding +6-8 °C above ambient. However, the authors could not decisively conclude whether this change was a direct effect of temperature or indirect effects due to, for example, effects on vegetation growth and phenology (Leblans et al., 2017), or an observed reduction in soil organic matter concentration, and soil texture (Poeplau et al., 2017; Verbrigghe et al., 2022). It is difficult to distinguish between potential drivers using community data alone, given that the relationship between specific taxonomic groups and ecological functions is largely unknown (Prosser, 2012).

A possible method for more precisely evaluating the direct effects of temperature on microbial communities is to directly characterize their physiological adaptation to temperature. It has been shown that soil bacterial communities adapt to the thermal environment, shifting measurable aspects of their aggregated temperature response (e.g. optimal [T_opt_], and theoretical minimal temperatures [T_min_] for growth) with changing temperature (Rinnan et al., 2009; Rousk et al., 2012; Bååth, 2018; Nottingham et al., 2019; Li et al., 2021). In an incubation study of alpine soils it was recently shown that such shifts in T_min_ occurred concurrently with a shift in bacterial community composition (Donhauser et al., 2020), but only when the imposed temperature treatment exceeded the measured T_opt_ of the *in situ* community. This result shows how targeted physiological assays can be used to complement community profiling methods to better understand the predominant mechanisms of microbial community variation (Hicks et al., 2021). A similar approach has been used to understand microbial community responses to soil pH changes (Pettersson and Bååth, 2013), and to link community composition and salt tolerance across a salinity gradient (Rath et al., 2019).

While community adaptation to temperature has been expressed in different ways (Bradford et al., 2008; Karhu et al., 2014; Dacal et al., 2019) the use of the apparent minimum temperature for growth (T_min_), estimated using the square root (also called the Ratkowsky) equation (Ratkowsky et al., 1982), provides an easily interpretable index (Pietikäinen et al., 2005; Bååth, 2018). In measurements taken on communities extracted from soils, T_min_ increases as a response to increasing soil temperature (Rinnan et al., 2009; Rousk et al., 2012; Bååth, 2018; Nottingham et al., 2019). This is also seen in pure culture studies, where differences in T_min_ can be used to differentiate between psychrophiles, mesophiles and thermophiles (Ratkowsky et al., 2005; Corkrey et al., 2016). Precise estimation of T_min_ requires a relatively large number of assay temperatures. If only relative changes in community adaptation to temperature are of interest, a more efficient method is to measure growth at two different assay temperatures, chosen to span a large part of the range between T_min_ and T_max_ (ideally making an interval that includes T_opt_), and to calculate a temperature sensitivity index (SI) as the log of the ratio of growth rates at high/low temperatures (Ranneklev and Bååth, 2001; Rinnan et al., 2009; Nottingham et al., 2019, 2021). A high value of SI indicates community adapted to relatively higher temperatures, while a low value reflects low-temperature adaptation. SI has earlier shown to correlate well with T_min_ (Rinnan et al., 2009, Nottingham et al., 2019).

In this study, we combined measures of temperature adaptation with taxonomic characterization of bacterial communities in order to test whether the previously observed shifts in bacterial community composition along the FORHOT geothermal warming gradient (Radujković et al., 2018) were related to community adaptation to temperature. We studied two temperature gradients, both with a gradient of ambient up to +40 °C, but with very different community composition (a forest and grassland soil). We hypothesized that in the part of the gradient with no difference in community composition there would be no temperature adaptation of the bacterial community, while a community change would be concomitant with increased SI indicating adaptation to higher temperatures. We also hypothesized that the same correlations would be found in the two different soils, despite having different original community composition. Thus, we expected temperature adaptation to be dependent on the temperature increase to the same degree in both soils.

## Materials and Methods

### Site description and sampling

The study area is the ForHot research site in the Hengil geothermal area, 40 km east of Reykjavik, Iceland. Mean annual temperature (MAT) is 5.2°C with the mean temperature of the coldest and warmest month (December and July) being −0.1°C and 12.2°C, respectively (Synoptic Station, Iceland Meteorological Office, 2016). There is usually no permanent snow cover during winter due to the mild oceanic climate, but the soil may freeze for at least 2 months during mid-winter. Sampling at the research site is concentrated in two vegetation types: an unmanaged grassland (in the area denoted “GN” in Sigurdsson et al., 2016) dominated by *Agrostis capillaris, Ranunculus acris* and *Equisetum pratense*; and a planted *Picea sitchensis* forest (“FN” in Sigurdsson et al., 2016), with no significant understorey vegetation. In May 2008 an earthquake shifted geothermal systems in the area surrounding the research site, resulting in hot groundwater warming the underlying bedrock and thus increasing the soil temperature in previously un-warmed soils. This resulted in transects with a gradient of increasing temperatures, from un-warmed controls to +40°C compared to the ambient temperature in the forest and grassland site. The area is described in more detail by Sigurdsson et al. (2016).

Soils were sampled in May 2012, that is 4 years after the beginning of warming. At each site four replicate transects were established, each consisting of 10 or 11 sampling plots located to cover to produce a set of temperatures on each transect. The target temperatures were: ambient, +1, + 2, +3, +4, +6, + 8, +20, +40 and plots were located at these temperatures using a handheld temperature probe inserted to 10cm depth. Subsequent monitoring of the soil temperature in each plot revealed some divergence from target temperatures, and for all analyses below we use the plot-level averages of temperature offset relative to ambient soils, measured on 4 – 10 different occasions in the period from October 2011 to July 2013 (see Appendix 1 for further discussion of temperature measurements). Note that although in the same area, these plots are not the same as those forming the permanent transects established in 2013 and described in several recent papers (Sigurdsson et al., 2016; Marañón-Jiménez et al., 2018; Radujković et al., 2018; Walker et al., 2018, 2020). For bacterial community analyses, three soil cores were taken at each plot with a 3cm diameter soil corer and the soil from 5 – 10 cm depth were retained for further analyses. Soil samples for community temperature adaptation measurements were taken 7 days later from the same sites using the same sampling method.

### DNA extraction, amplification and sequencing

Soil samples were stored at 4°C within two hours of sampling, and processed in the following 48 hours. Soils were sieved and mixed, and DNA extracted using MoBio PowerSoil DNA kit following the manufacturer’s protocol. DNA samples were checked for quality and quantity using a Nanodrop spectrophotometer and prepared for Illumina sequencing of the V3 region of the bacterial 16S rRNA gene using the primers in Bartram et al. (2011) following the procedures described in Weedon et al. (2017). Libraries were sequenced on an Illumina MiSeq using 2x 150 cycle paired-end sequencing (V2 chemistry) at the VUmc Clinical Genetics sequencing facility (Amsterdam, The Netherlands).

### Bioinformatics

Initial sequence processing and OTU clustering was performed using the USEARCH software (Edgar, 2013). Paired-end sequences were assembled with maximum of 3 mismatches allowed in the overlapping region (77% of raw-reads retained). This was followed by quality filtering with maximum expected errors set at 0.05 which removed an additional 23% of the successfully merged reads. Operational taxonomic units (OTUs) were then defined using the UPARSE algorithm with 97% minimum similarity (Edgar, 2013), after removing all singleton reads. Chimeric sequences were removed with UCHIME (Edgar et al., 2011). A set containing representative sequences for each OTU was aligned using PyNAST (J Gregory Caporaso et al., 2010) using as a reference alignment the Green Genes version 13_8 (DeSantis et al., 2006) ‘core-set’ as distributed with QIIME version 1.7.0 (J. G. Caporaso et al., 2010). Sequences belonging to OTUs that failed to align with at least 75% sequence similarity, were most likely chimerical sequences or sequencing errors, and were removed from the dataset (178 OTUs representing 0.6% of successfully assembled reads). All original reads were mapped back onto the resulting OTUs to produce an OTU table. For phylogenetic distance measures we generated a phylogenetic tree based on the aligned representative set using FastTree (Price et al., 2009). Lastly, we assigned all OTUs to a taxonomic classification using the Ribosomal Database Project Bayesian classifier (Wang et al., 2007) with a threshold minimum confidence of 80%. Raw sequences will be deposited in the NCBI Sequence Read Archive.

### Community temperature adaptation measurements

Soil samples were stored at 17°C until analysed (within 2 months). The higher temperature used (not the more commonly used 4°C) was in order to not affect temperature adaptation in the soils with highest temperature. Still, this temperature has been shown not to affect temperature adaptation during this time period (Bárcenas-Moreno et al., 2009; Birgander et al., 2013). Adaptation of the bacterial community to temperature was measured using a growth based index, where bacterial growth was estimated as leucine (Leu) incorporation (Bååth, 1994; Bååth et al., 2001). Bacterial growth was determined at two temperatures, 40°C and 4°C, and the log ratio (denoted log (growth at 40°C/4°C) was used as sensitivity index (SI) of temperature adaptation of the bacterial community. A high value indicates adaptation to high temperatures, a low value to low temperature conditions (Rinnan et al., 2009; Nottingham et al., 2019). The index correlates to the more commonly used T_min_ (minimum temperature for growth) to express temperature adaptation (Nottingham et al., 2019). Due to the large range of in *situ* temperatures, a larger difference in the high and low incubation temperature was used compared to earlier studies.

For each replicate, bacteria were extracted from soil (1 g soil in 25 mL water) by shaking, followed by a low speed centrifugation (1000 x g for 10 min). The bacterial suspension was then distributed in microcentrifugation vials (1.5 ml in each) and ^3^H-Leu 37 MBq mL^−1^ and 5.74 TBq mmol^−1^, Perkin Elmer, USA) and unlabeled Leu (final concentration of 275 nmol L^−1^) was added after 30 min at the chosen incubation time. Growth was terminated by adding trichloroacetic acid after incubating for 2.5h at 40°C and 24h at 4°C. Incubation times were chosen in order to achieve more similar total Leu incorporation at the two temperatures. Due to logistic reasons measurements at the different temperatures were made at different days. Subsequent washing steps and measurement of radioactivity were performed following the procedure described by Bååth et al. (2001).

Control soils were also subjected to growth analysis at a range of temperatures (5°C intervals between 0°C and 50°C) in order to determine original T_min_ and T_opt_ for bacterial growth. T_min_ was calculated by the Ratkowsky equation (Ratkowsky et al., 1982) on square root transformed data at temperatures below T_opt_ (≤25°C, see Fig. 1). A relationship between T_min_ and the log ratio growth at 35°C/0°C was found in a gradient spanning 20°C in mean annual temperature (Nottingham et al., 2019). This was used together with a conversion factor from the growth ratio 40°C/4°C to 35°C/0°C (unpublished) to calculate corresponding values of T_min_ from growth ratios 40°C/4°C.

**Figure 1:**
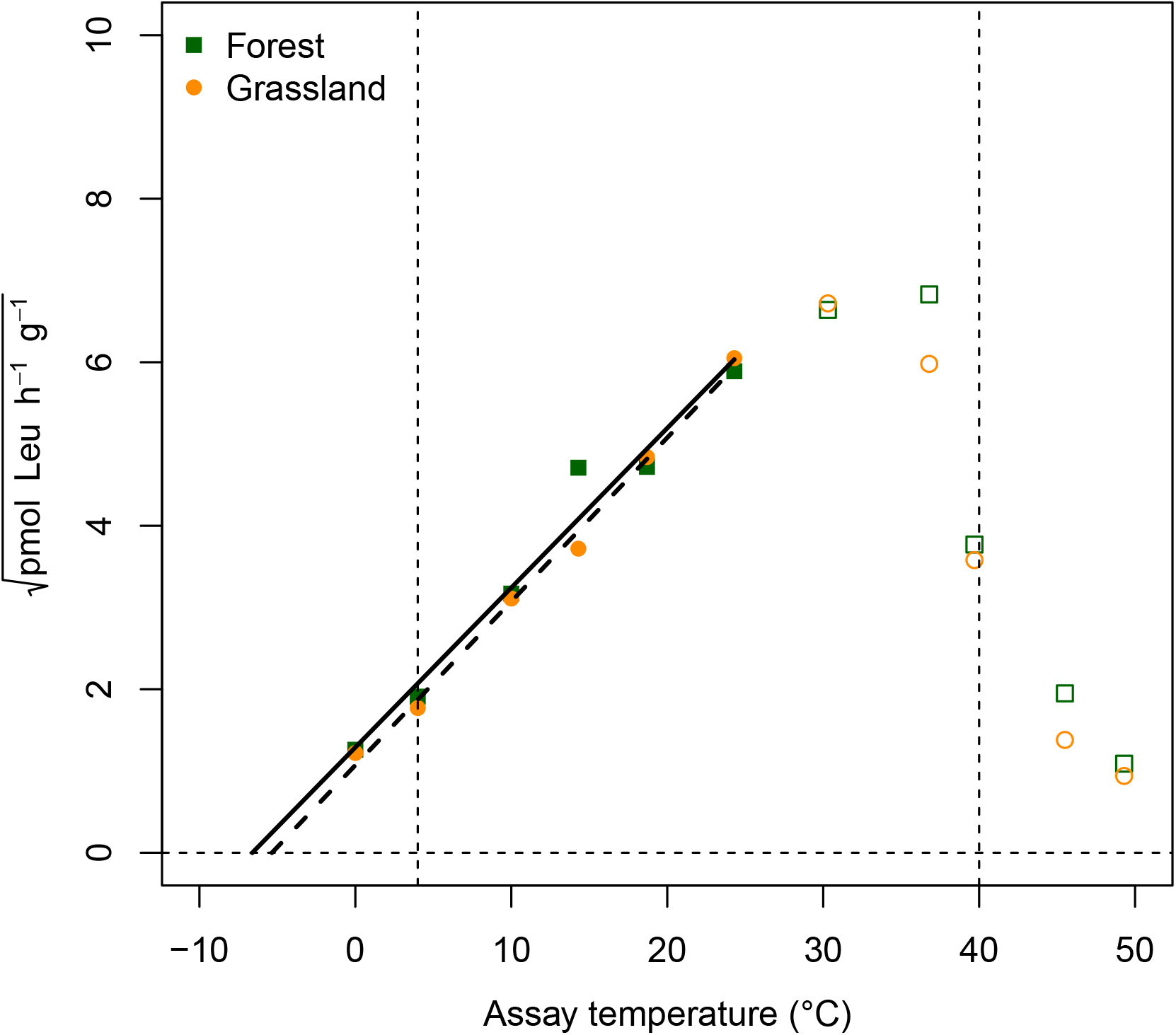
Temperature dependence of growth (Leu uptake) from bacterial communities sampled from soils with ambient temperatures (MAT 5.2°C) in the grassland (GN, squares and solid line) and forest (FN, circles and dashed line) habitats. The data are plotted using square root transformation, with filled symbols included in calculations using the square root (Ratkowsky) equation. *T_min_* was −6.6°C and −5.4°C in GN and FN, respectively. Thin vertical lines indicate temperatures used for the Sensitivity index (SI, log(growth at 40°C/at 4°C).

### Statistics

Alpha diversity (Shannon index) was computed for each bacterial community sample. Pairwise weighted Unifrac distances (Lozupone et al., 2006) were computed for all sample combinations and visualized using principle coordinates analysis (PCoA). The first PCoA axis (explaining 25% of the dataset variation) was subsequently used as a univariate proxy of community composition. The significance and relative magnitude of site and temperature elevation effects on total community composition was analysed with a permutational multivariate analysis of variance (Anderson, 2001) on the Unifrac distance matrix.

Alpha diversity indices, PCoA scores and SI were each modelled as a function of temperature elevation separately for each site. Initial visualization indicated a step-wise pattern for all responses at both sites. This was further investigated by fitting regression trees to each dataset and comparing the resulting break-point model with normal linear regression using AIC values. Given that there was some uncertainty as to the temperature elevations at each plot (see Appendix 1) we used a parametric bootstrap approach to estimate the uncertainty associated with the breakpoint in the tree regression for each response x site combination. Bootstrap distributions of the breakpoint were generated by generating 1000 sets of temperature points created by adding Gaussian distributed noise to each of the temperature values, and refitting the regression tree model. The standard deviations of the added noise were based on the degree of confidence of each interpolated temperature elevation (ranging from 0.5 to 2.5 °C). To directly test for a relation between community temperature adaptation and composition, separate linear regressions were performed between PCoA1 scores and SI for each site.

To identify bacterial OTUs response to warming we performed a differential abundance analysis using the ANCOMBC R-package (Lin and Peddada, 2020). For both sites, we filtered samples to a minimal sequencing depth of 3750 reads, and filtered OTUs on an abundance above 0.001 % and minimal occurrence in 3 samples. We ran ANCOMBC for soil bacterial community data of both sites separately, using the increase in soil temperature as independent variable and transect as a covariate, with Bonferroni method as for false discovery rate correction.

## Results

### Community adaptation to temperature

Bacterial growth in ambient soils from both sites closely followed the Ratkowsky model with the square root of growth increasing linearly with temperature below T_opt_ (Fig. 1). T_min_ of bacterial growth was similar in the two soils, −6.6°C and −5.4°C in the grassland and forest, respectively. T_opt_ was not specifically determined, but appeared to be around 30°C. Above T_opt_ bacterial growth decreased rapidly with increasing temperature. In the ambient soils SI (log ratio of growth at 40 and 4°C, see Fig. 1) was 0.59 and 0.61 in the grassland and forest, respectively.

### Bacterial community composition

In ambient soils the bacterial communities of both the grassland and forest sites were dominated by OTUs assigned to the phyla Proteobacteria (35% of reads in the forest site, 31% in the grassland), Actinobacteria (15% and 21%) and Acidobacteria (26% and 20%). Other phyla contributing between 1 and 8% of reads were Chloroflexi, Firmicutes, Bacteroidetes, Verrucomicrobia, Nitrospirae, and Gemmatimonadetes. Bacterial community composition was significantly different between the two vegetation types (Weighted Unifrac PERMANOVA *P* = 0.004, R^2^ = 0.27, *n* = 13).

### Soil temperature effects on bacterial community composition

The bacterial community was significantly affected by increasing soil temperatures at both sites. This was visible at the level of OTU diversity (Fig. 2, breakpoint regression P < 0.05), overall community composition (Fig. 3, weighted Unifrac PERMANOVA temperature effect: P = 0.001, R^2^ = 0.12), and the a large number of differentially abundant OTUs (Fig. 4; ANCOMBC; P<0.05).

**Figure 2:**
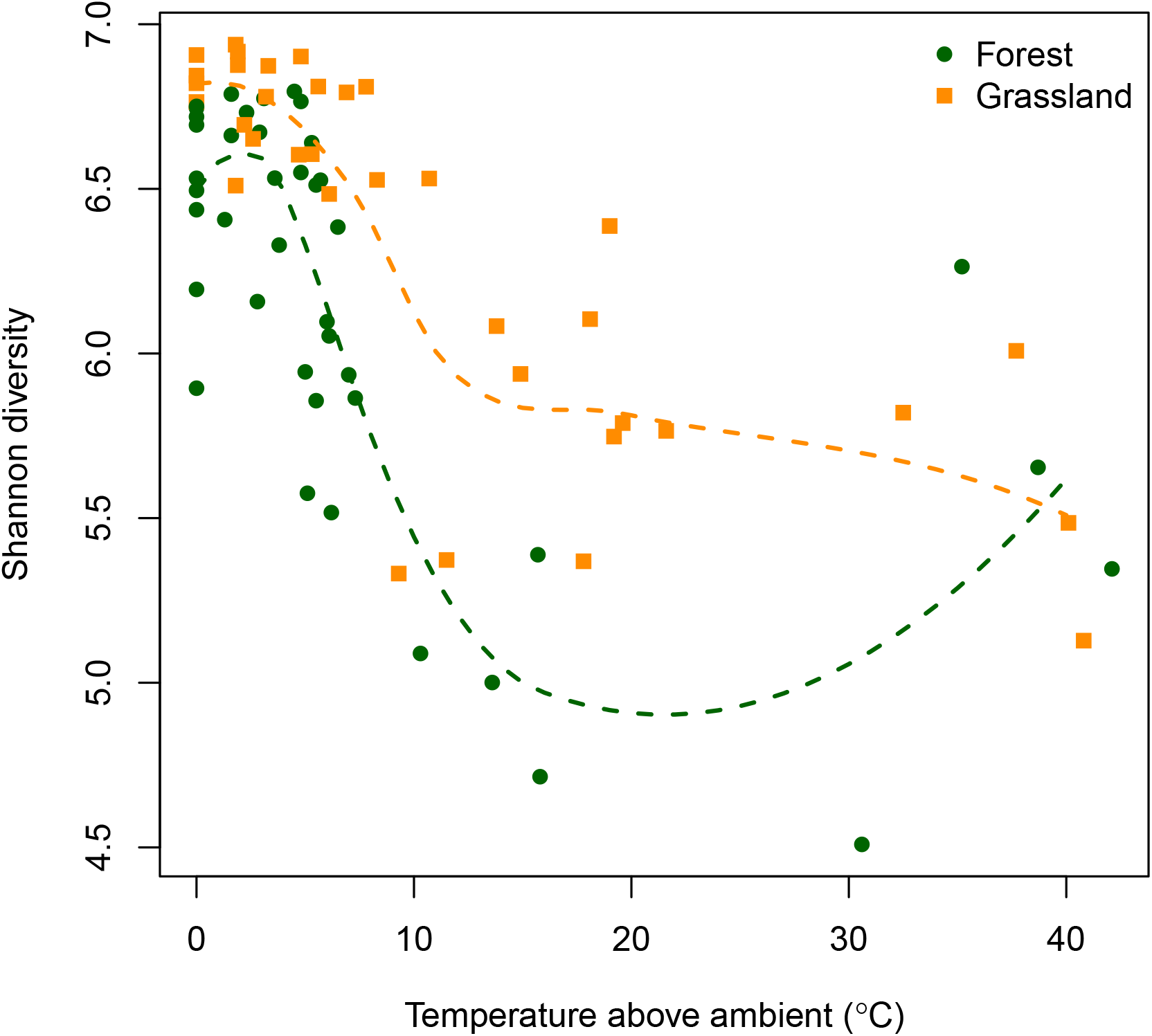
Alpha diversity of the bacterial community (Shannon index) versus soil temperature above ambient (MAT 5.2°C) along two geothermal soil temperature gradients (grassland, GN, yellow squares and forest, FN, green circles). Dashed lines represent a loess smoothing function.

**Figure 3:**
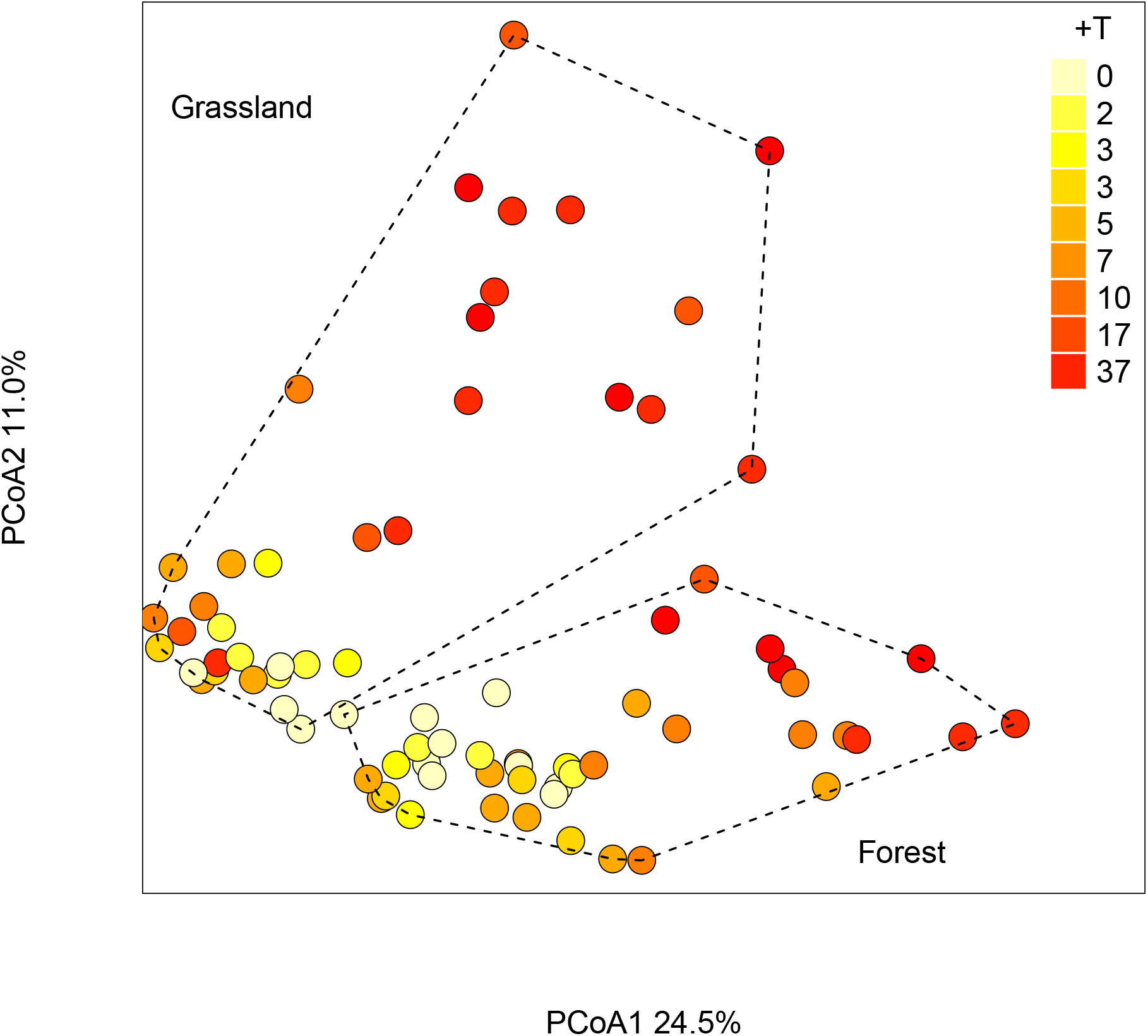
Principal coordinates analysis ordination (PCoA) of bacterial community profiles from two geothermal soil temperature gradients (grassland, GN, and forest, FN). Ordinations are based on weighted-Unifrac distances computed from 16S rRNA gene amplicon data and colour coded according to temperature elevation (in °C) above ambient soils (MAT 5.2°C)

**Figure 4:**
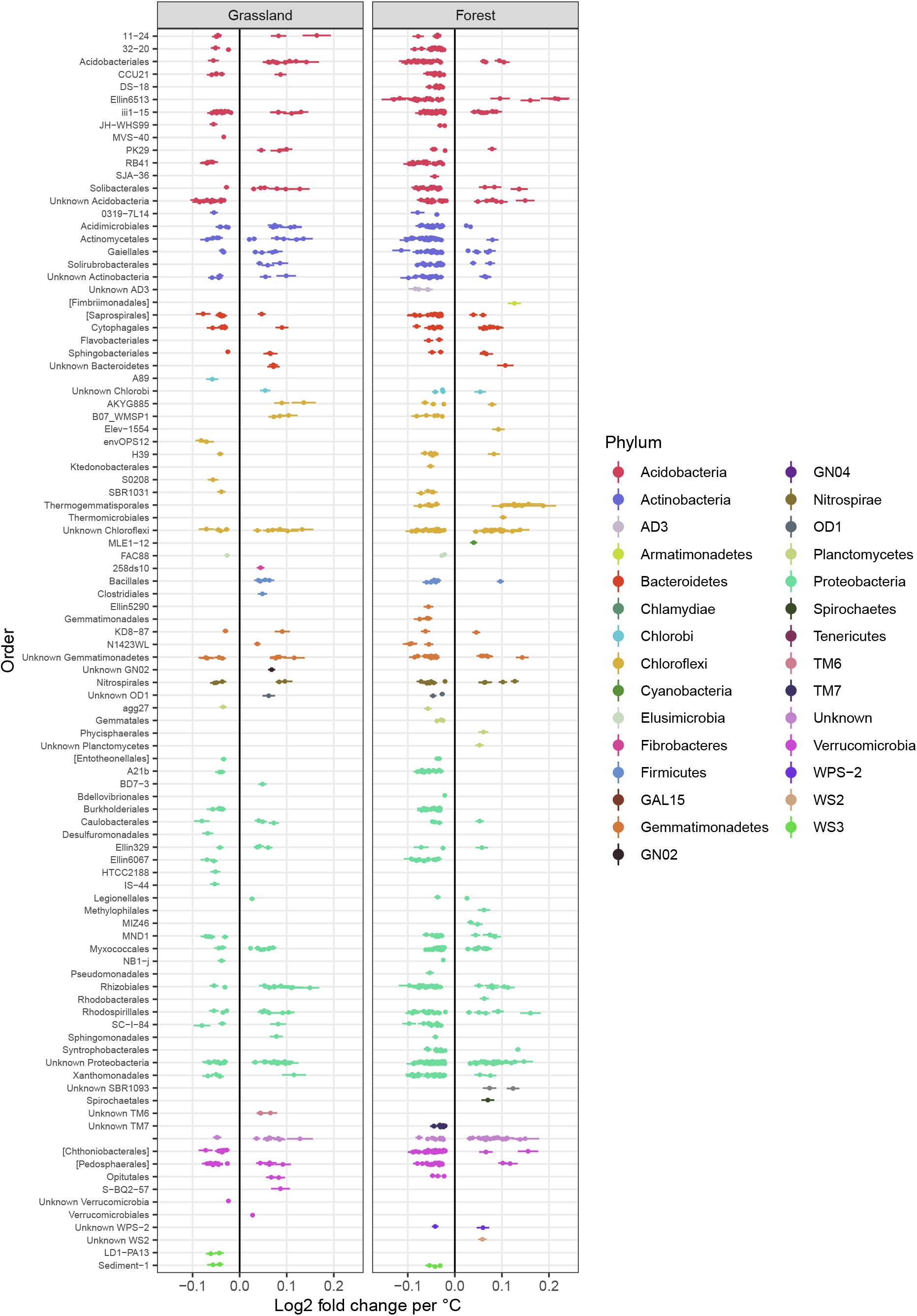
Abundance shiftsof the OTUs that were differentially abundant (P<0.05) across the warming gradient for both Forest and Grassland, expressed in log_2_-fold changes and grouped by Order rank on the y-axis. Colors indicate Phylum rank.

Alpha diversity decreased from around 6.5 to 7 in both ambient sites to between 5 and 6 in the warmer soils (Fig. 3). This decrease in diversity was paralleled with a changing community composition, with the forest soil mainly changing along PCoA1 (explaining 24.5% of the variation), while temperature effects was detected in both PCoA1 and PCoA2 (11.0% of the variation) for the grassland community composition (Fig. 3). The temperature effect was of roughly the same magnitude as site differences in explaining variation in the bacterial community composition (weighted Unifrac PERMANOVA: temperature effect: P = 0.001, R^2^ = 0.12, site effect: P = 0.001, R^2^ = 0.15).

The overall shifts in community composition were made up of changes in relative abundance of a large number of OTUs. In total, 849 out of 2926 OTUs for the forest site and 303 out of 3120 OTUs for the grassland site showed differential abundance across the soil warming gradient. Only 104 OTUs were differentially abundant in both sites, of which most OTUs belonged to the phyla Proteobacteria, Acidobacteria, Actinobacteria, and Verrucomicrobia (Fig. 4, Table 1). In the forests soils, the 186 OTUs that increased in abundance went from contributing 0.7 % of the total reads in the ambient soil samples to 49.3 % in the soils warmed above 6 °C. In total 663 OTUs decreased in abundance significantly, reducing the abundance of the total group from 60.2 to 17.3% of the total reads. In grassland soils, 142 OTUs increased in abundance from 3.3 % of the total reads in the ambient soil to 25.5 % in the soils warmed above 6 °C. In total 161 OTUs decreased in abundance significantly, reducing in their contribution to total reads from 13.7 to 3%.

**Table 1.**
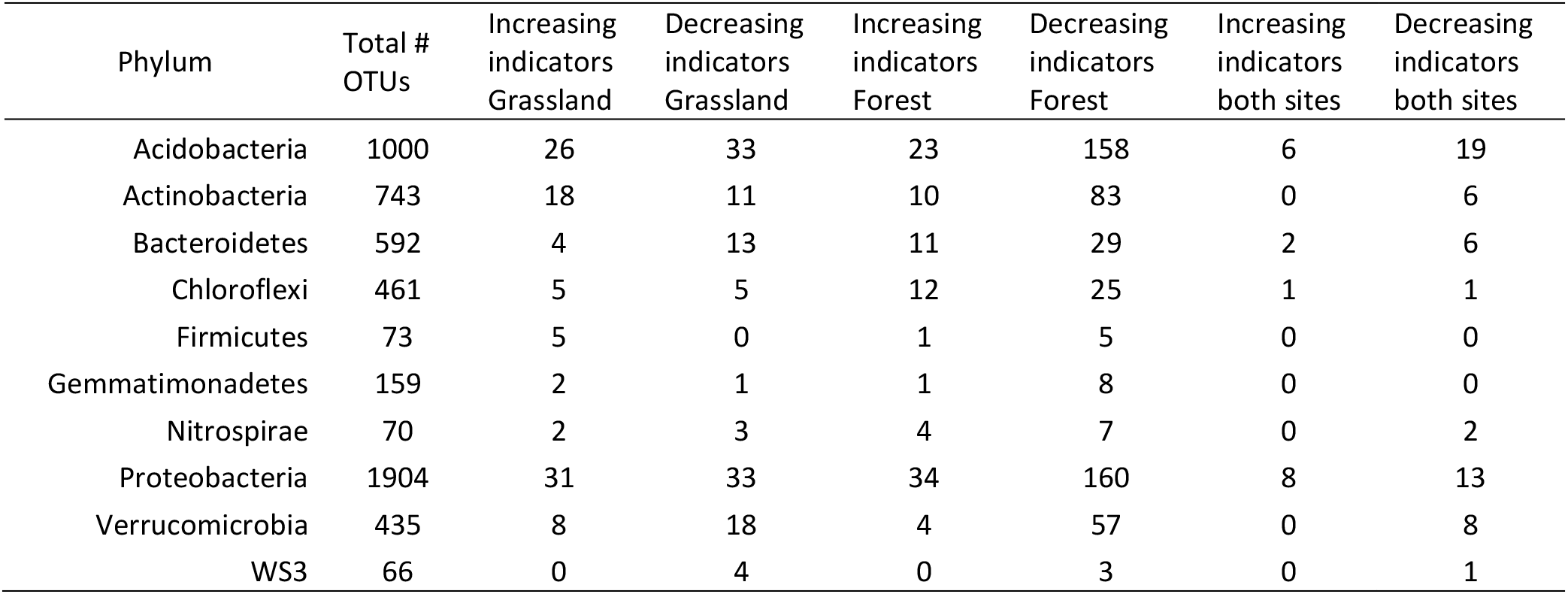
Distribution of bacterial OTUs across phyla (only top 10 most abundant phyla shown). Total number of OTUs as well as numbers observed to significantly differ (positively or negatively) between samples above or below 6°C in one or both sampling sites. Significant differences were calculated using ANCOM-BC (*P* < 0.05, after Bonferroni false discovery correction).

| Phylum | Total # OTUs | Increasing indicators Grassland | Decreasing indicators Grassland | Increasing indicators Forest | Decreasing indicators Forest | Increasing indicators both sites | Decreasing indicators both sites |
| --- | --- | --- | --- | --- | --- | --- | --- |
| Acidobacteria | 1000 | 26 | 33 | 23 | 158 | 6 | 19 |
| Actinobacteria | 743 | 18 | 11 | 10 | 83 | 0 | 6 |
| Bacteroidetes | 592 | 4 | 13 | 11 | 29 | 2 | 6 |
| Chloroflexi | 461 | 5 | 5 | 12 | 25 | 1 | 1 |
| Firmicutes | 73 | 5 | 0 | 1 | 5 | 0 | 0 |
| Gemmatimonadetes | 159 | 2 | 1 | 1 | 8 | 0 | 0 |
| Nitrospirae | 70 | 2 | 3 | 4 | 7 | 0 | 2 |
| Proteobacteria | 1904 | 31 | 33 | 34 | 160 | 8 | 13 |
| Verrucomicrobia | 435 | 8 | 18 | 4 | 57 | 0 | 8 |
| WS3 | 66 | 0 | 4 | 0 | 3 | 0 | 1 |

### Soil temperature effects on bacterial community adaptation to temperature

Increasing soil temperatures resulted in increased SI in both the forest and the grassland (Fig. 5), indicating growth adaptation to higher temperatures of the bacterial community along the geothermal gradient at both sites. SI increased from around 0.6 in both ambient sites to around more than 1.5 in the warmest plots. Recalculating these changes in SI to approximate changes in T_min_ resulted in an increase from a T_min_ of around −6°C in ambient sites to around − 1 °C in the sites with >+15°C above ambient temperatures (MAT increasing from +5°C to >20°C). Thus, T_min_ increased around 5°C with an increase in MAT of >15°C.

**Figure 5:**
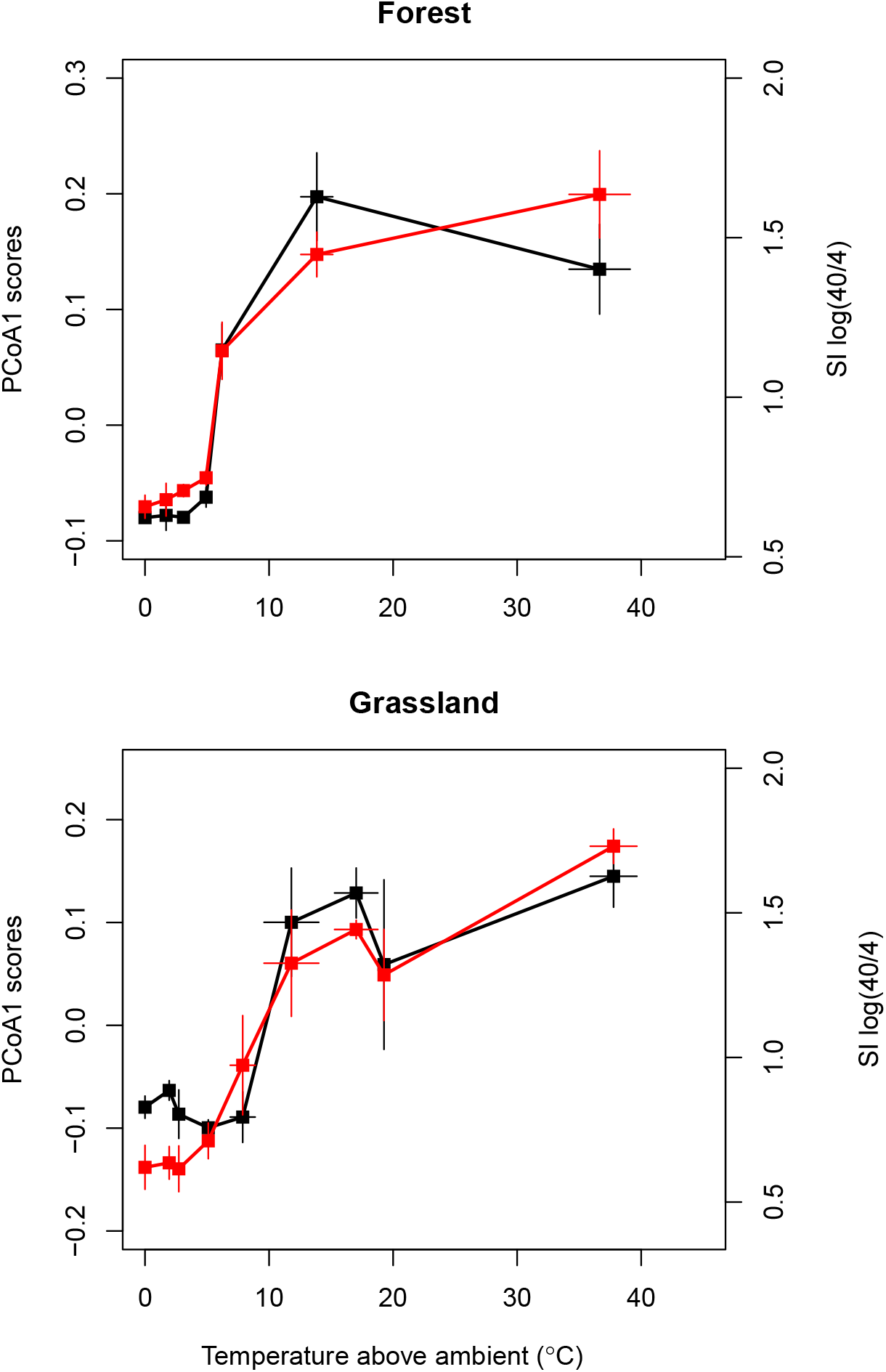
Relationships between bacterial community composition and community adaptation to temperature along soil temperature gradients. Bacterial community expressed as axis scores of PCoA ordinations of UniFrac distances (black points and line) and community growth adaptation as SI, (log growth at 40°C/at 4°C) (red points and line) as a function of soil temperature elevation above ambient. For visualization, lines join mean values (large points) computed for groups of samples with similar temperature elevations; error bars are standard errors of the mean for both the response and the soil temperature elevation (n = 3-9 samples per group). Actual statistical modelling (see Table 1, Figure 6) was performed on plot-level observations.

### Comparing temperature effects on bacterial community composition and growth adaptation

There appeared not to be a gradual effect of temperature along the geothermal gradient on the soil microbial community. Instead, for all of the aforementioned measures (community profile (PCoA1), Shannon diversity, growth adaptation to temperature (SI)), with the exception of SI in the grassland site, a break point model as a function of soil temperature was a closer fit to the observed data than either simple linear regression or a null model (Table 2, all models P < 0.05). There was therefore a threshold temperature, below which there appeared to be no effect on the bacterial community. Median threshold temperature was 7.5, 6.1 and 8.6°C for community profile, SI and Shannon Diversity in the forest site, with corresponding values for the grassland being 9.5, 9.0 and 9.0°C (Fig. 6). However, when accounting for uncertainty in soil temperature measurements, the range of estimated threshold temperatures spanned 5 to 12 ° C (end points of bootstrap 95% confidence intervals) with relatively more uncertainty at the grassland site.

**Table 2.** Model selection metrics (AIC) comparing linear regression and single breakpoint stepwise function models for each of community profile (PCoA axis 1 scores), temperature sensitivity index (SI, log (growth at 40°C/4 °C)), and bacterial alpha diversity (Shannon) for both sites.

|  |  | Linear Model | Breakpoint model |
| --- | --- | --- | --- |
| | | ($k = 3$) | ($k = 4$) |
| Forest | Community profile ($n = 41$) | -83 | **-101** |
| | SI ($n = 35$) | 1.8 | **-7.6** |
| | Alpha diversity ($n = 41$) | 63 | **47** |
| Grassland | Community profile ($n = 35$) | -72 | **-81** |
| | SI ($n = 42$) | **2.9** | 4.1 |
| | Alpha diversity ($n = 35$) | 31 | **15** |

**Figure 6:**
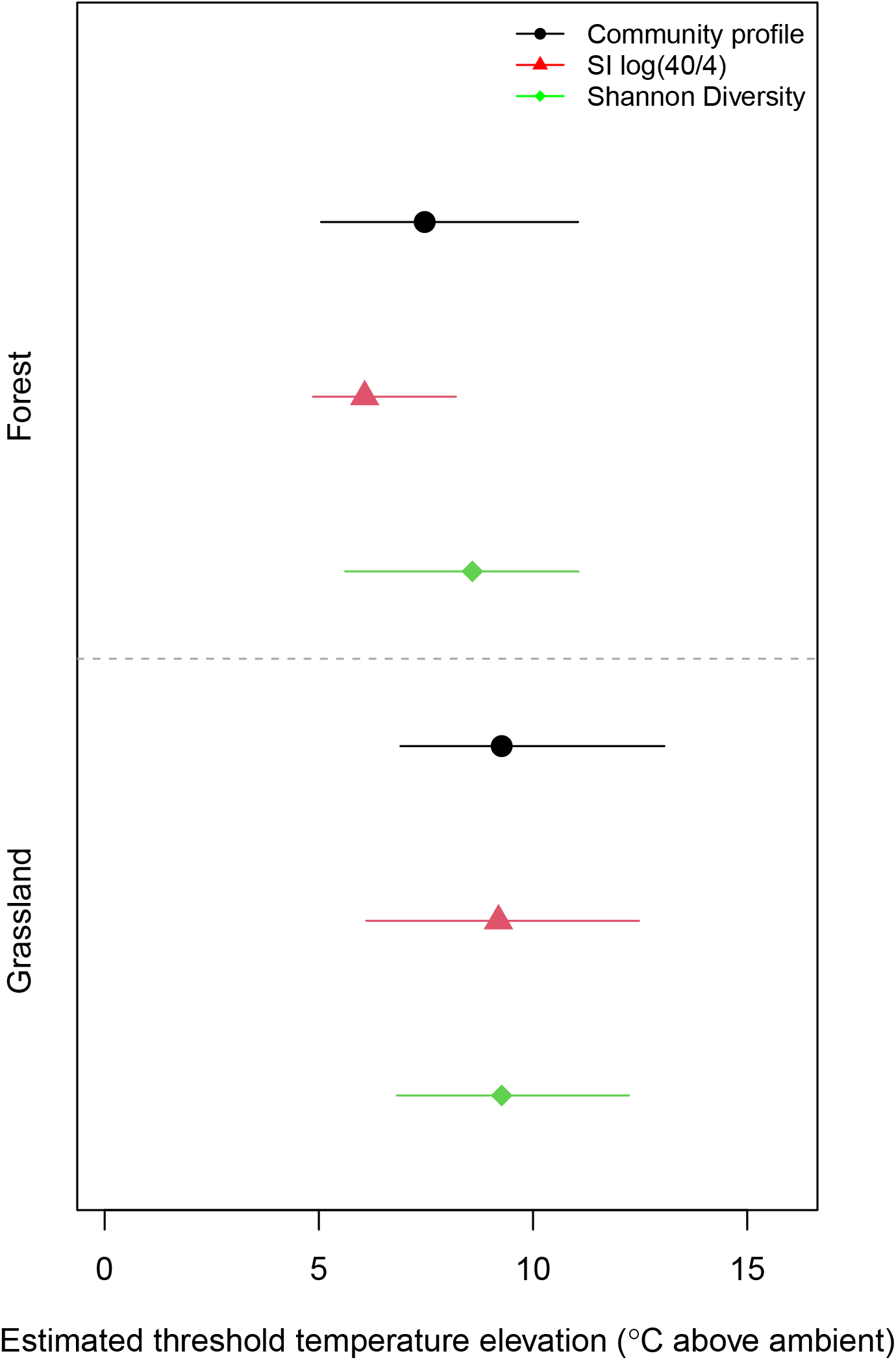
Estimates of temperature thresholds for significant changes and associated uncertainties for bacterial community composition (PCoA1 scores, see Fig. 3), growth adaptation to temperature as a Sensitivity Index (SI, log growth at 40°C/at 4°C), and Shannon diversity at two geothermal soil temperature gradients (grassland, GN, and forest, FN). Distributions were generated using parametric bootstrapping of temperature data, based on computed uncertainties in temperature estimates (for methology see Supplements). Bars and points correspond to the 95% confidence interval and median, respectively, of bootstrap distributions of the breakpoint in tree regression models.

As noted above, the observed shifts in bacterial community composition and SI with temperature followed very similar dynamics in both sites (Fig. 5). Accordingly, there were significant linear correlations between these two measures at both sites (R^2^=0.78 and 0.74, for forest and grassland, respectively (Fig. 7).

**Figure 7:**
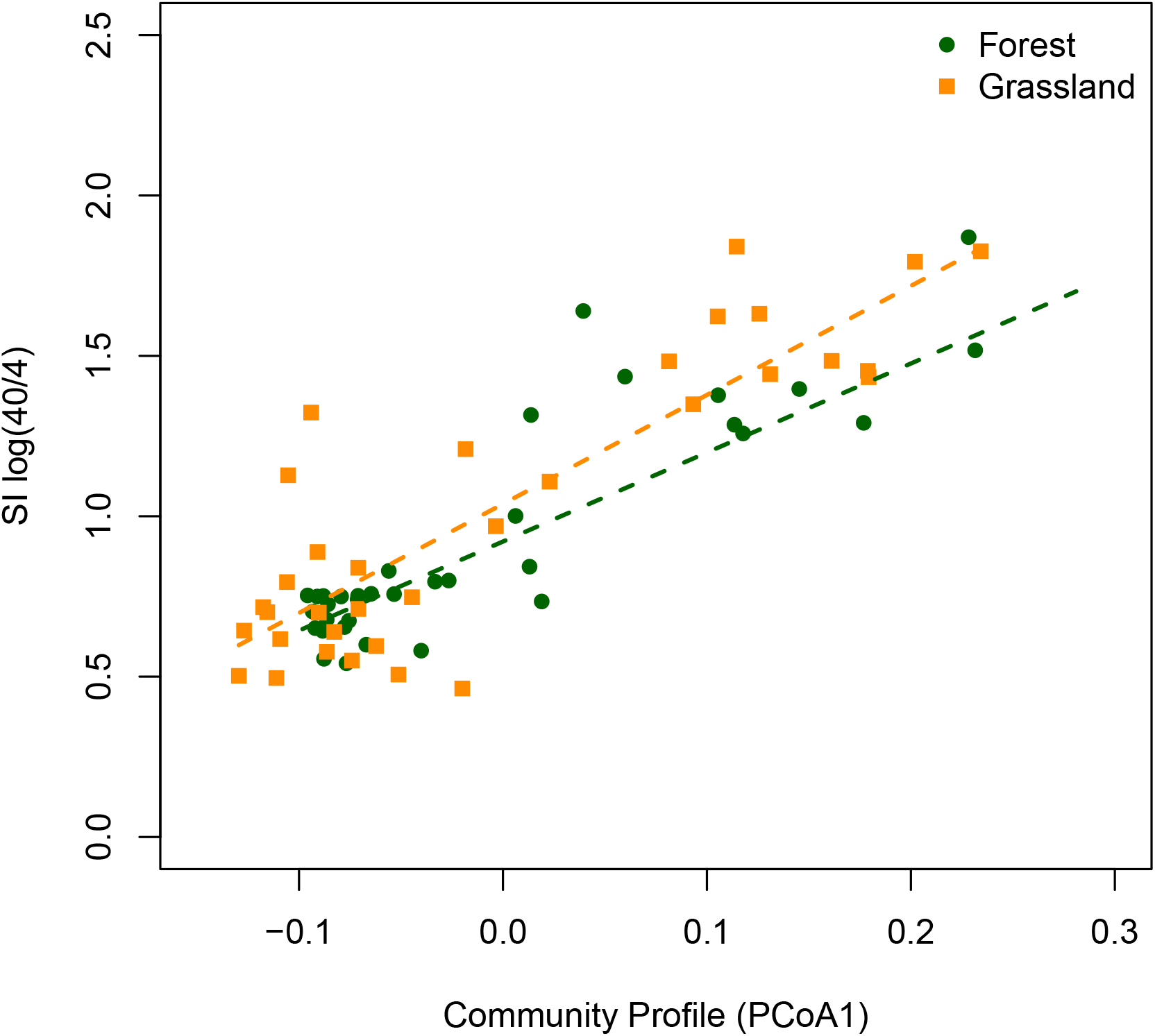
Relationships between bacterial community composition (PCoA1 scores, see Fig. 3) and growth adaptation to temperature as a Sensitivity Index (SI, log growth at 40°C/at 4°C) for samples along two geothermal soil temperature gradients (grassland, GN, orange squares, and forest, FN, green circles). Lines represent linear regressions, in both cases P < 0.05, *R*^2^ = 0.78 (forest), 0.74 (grassland).

## Discussion

In this study we characterized bacterial community composition and physiological temperature adaptation of sub-arctic soils sampled along a 40 °C soil warming gradient under two vegetation types. We confirmed previous observations that bacterial community profiles only change significantly relative to ambient conditions when subject to more than approximately 6-9 °C warming (Radujković et al., 2018; Walker et al., 2018). Crucially, we provide the first evidence that this community shift is driven by direct responses to temperature, by showing that community change coincided closely with increases in temperature adaptation of the community. This overlapping of community trait and compositional measures was observed in both grassland and forest soils, even though each environment had distinct bacterial community profiles at both ambient and warmed conditions.

### Bacterial community composition responses to warming

The bacterial community composition in both grassland and forest soils shifted in response to increasing soil temperatures (Fig. 3). The shift was characterized by a decrease in alpha diversity, which was demonstrated by more OTUs declining in abundance over the warming gradient than OTUs increasing in abundance (Table 1). This is contrary to studies finding increasing diversity with increasing temperature, for example in elevation gradients (Nottingham et al., 2018; Ji et al., 2022). This suggests different effects of temperature in response to long-term (natural temperature gradients) versus short-term warming (4 years in the present temperature gradient).

The warm responders made up a small fraction of the bacterial community (0.6% in forest and 3.3% in grassland) in the ambient soil but increased to 48.7 and 25.5% of the total community above 6°C warming in the forest and grassland, respectively. A similar (partial) community turnover was also observed in an incubation of alpine soils, where increasing temperatures lead to a community dominated by presumably warm-adapted taxa (Donhauser et al., 2020). Community-level responses to warming, either experimentally imposed, or measured along climatic gradients, have been observed for bacteria in a number of soil environments (Yergeau and Kowalchuk, 2008; DeAngelis et al., 2015; Oliverio et al., 2017; Monteux et al., 2018; Radujković et al., 2018), although they are not always present (Weedon et al., 2017). The responding OTUs appeared to be fairly evenly distributed amongst the major phyla, and within each phylum there were OTUs that both increased and decreased with soil warming. This emphasizes that temperature niche seems to be largely decoupled from high-level taxonomic identity (Oliverio et al., 2017; Radujković et al., 2018) supporting the idea that temperature preference of bacteria is a shallowly conserved trait (Martiny et al., 2015).

### Bacterial community adaptation to temperature

The bacterial community adaptation to temperature, which we estimated based on the log ratio of bacterial growth rates measured at 40 and 4 °C (SI), was relatively constant along the warming gradient up to between 6 and 9°C before abruptly increasing. Most previous studies of the relationship between soil temperature and temperature adaptation of bacterial community growth have expressed the results in terms of T_min_ (the theoretical minimum temperature for growth) derived from fitting the Ratkowsky square root model to temperature-growth data (Bååth, 2018). Nottingham *et al*. (2019) showed that there was a close correlation between T_min_ and SI, meaning that not only is SI a good indicator of temperature adaptation, but our observed SI shift would be approximately equivalent to an increase in T_min_ from around −6°C to between −1 and −2 °C at temperatures >15°C above ambient - an increase of 0.27 to 0.33°C per 1°C increase in temperature. A T_min_ of −6°C with a MAT of +5°C is within the expected range for this subpolar oceanic climate (Bååth, 2018). The increase due to the soil warming is also comparable to earlier estimations based on climatic gradients (Rinnan et al., 2009; Nottingham et al., 2019), where an increase of 0.2 to 0.5°C per degree increase in MAT is typically found. Higher values (around 0.8 per 1°C warming) have also been reported from incubation studies (Birgander et al., 2013; Donhauser et al., 2020), whereas field warming studies have tended to estimate smaller or even absent effects (Rinnan et al., 2011; Rousk et al., 2012). Still, the increase in T_min_ per degree increase in MAT appears always to be less than one. Bååth (2018) suggested, based on the studies above, a tentative value for a T_min_ increase of 0.3°C/degree increase in MAT, which is well in accordance with our results.

An alternative to SI, although less sensitive, is to compare Q10 values (the fold change per 10°C warming of a biological process) measured over a defined interval. For example, Q10 of soil respiration increases at higher temperature conditions and decreases at cooler temperatures (Kirschbaum, 1995). Previous studies of respiration at the FORHOT grassland site have used this approach (Marañón-Jiménez et al., 2018; Walker et al., 2018), reporting the same temperature sensitivity in warmed areas compared to control sites. This implies no community adaptation to increasing temperature in contradiction of our results from growth assays. In the case of the results from Walker et al. (2018) only soils with warming up to 6°C above ambient temperature were included, that is, their results are compatible with the present study, since this is similar or less than our estimated threshold temperature. Marañón-Jiménez et al. (2018) studied respiration of grasslands soils with warming levels up to 15.9°C without any significant changes in the apparent temperature sensitivity of microbial respiration. One reason for this different result may be due to varying sensitivity of temperature adaptation for soil respiration as opposed to bacterial growth. However, Bååth (2018) compiled data on both bacterial growth and respiration and found that these processes appeared to have comparable T_min_ in soils globally, and therefore presumably similar responses to warming. Li et al. (2021) also found that T_min_ for soil respiration correlated to MAT in a large set of soils covering alpine to tropical soils. An alternative explanation for the discrepancy could be the relative large variations in the respiration data, as well as the use of fewer soils sampled from areas with temperatures at >8°C above ambient (only 2 areas in one site compared to our two sites with 3-5 areas each). Indeed, a closer inspection of their results (Table 3 in Marañón-Jiménez et al. (2018)) reveals that sites with Amb+<8°C had Q10 values of 2.29 to 2.83, while the two sites with Amb+>8°C both had higher Q10 for respiration, 3.09 and 4.77, suggesting a lack of statistical power may have led to the conclusion of no change in temperature sensitivity over the warming gradient. Instead, a threshold in temperature adaptation similar to described in the present study may also be present for soil respiration.

### Mechanisms behind threshold dynamics

For all community composition measures the change with temperature was abrupt rather than gradual, with the critical warming level triggering the shift estimated to be between approximately 6 and 9 °C above ambient conditions, depending on the vegetation type and community measure. The clear evidence of a threshold dynamic in the response of community composition at both sites to increasing soil temperature confirmed previous observations from this geothermic area (Radujković et al., 2018; De Jonge et al., 2019). However, our study provides the first evidence that this shift is accompanied by a change in the temperature adaptation of the bacterial community. We do not know whether the abrupt shift of bacterial communities we observed is unique to our study site, or reflects a warming response in a wider range of soils and ecological contexts. If the pattern is indeed more common, it may explain some of the divergence in results of warming experiments. Studies that show non-effects of warming on soil bacterial communities may have involved experimental warming treatments that fail to surpass the critical threshold temperature. Given that we estimate a threshold warming of > + 6 °C, this would explain the relatively minor effects of open top chamber treatments (typical warming effects 0.5 - 3 °C, Marion et al., 1997) and even soil heating cables (typical warming effects ~ 5 °C, Rustad et al., 2001).

Threshold dynamics in community changes with warming could arise as the results of an ecophysiological ‘tipping point’ that is common to bacterial communities. It has been suggested that warming that is close to or exceeds the community aggregated T_opt_ for growth represents such a limit (Bárcenas-Moreno et al., 2009; Donhauser et al., 2020), but the result from the present study is not in accordance with this, since even our highest estimate of the threshold warming represents a soil MAT of approximately 17 °C, compared to a community T_opt_ of about 30 °C in ambient soils (Fig. 1). Furthermore, changes in T_min_ has been found in gradient studies at low temperatures, where changes in MAT is well below T_opt_ for bacterial growth (Rinnan et al., 2009; Nottingham et al., 2019) An alternative explanation for the shape of the response are time lag effects; that is, changes take time, especially at low temperatures. It is possible that given enough time all communities reach a compositional and physiological “equilibrium” state relative to soil temperature. However, if the rate of this process of equilibration is itself driven by temperature, then warmer soils will reach the new steady state more rapidly than areas with more modest warming (Pettersson and Bååth, 2003). In our case, the threshold temperature is an increase from a low ambient MAT (5.2°C). This means that even ambient +6°C of warming only has a MAT of around +11°C. This interpretation is supported by the observation from a reciprocal transplant study over an altitudinal gradient that showed slower (decade-scale) temperature adaptation to cooler environments compared to adaptation to warmer environments (Nottingham et al., 2021). However, a comparison of grassland sites with different warming durations (50+ years vs 4 years, the same site as the current study) did not produce any evidence to support this time lag interpretation (Radujković et al., 2018).

### Implications and further research

Regardless of the underlying mechanism responsible for the observed threshold dynamics, the observation that a major shift in community composition occurred at the same level of warming as a similarly abrupt change in SI provides strong evidence that shifts in the composition of the community was driven by direct temperature effects. A large variety of soil and vegetation parameters have also changed along this geothermal gradient (Sigurdsson et al., 2016; Walker et al., 2020), implying that at least some of observed changes in bacterial community could be indirect effects. However, the close relationship between community composition and SI would be unlikely if drivers such as substrate, pH, or soil texture were the dominant control. Even so, given the inevitable confounding of potential drivers in natural settings, controlled experiments that manipulate temperature, while holding other important driving factors constant, are still needed to confirm the predominance of direct temperature adaptation effects in determining responses of bacterial community composition to warming.

Oliverio et al. (2017) compiled data on bacterial taxa that could be used as indicator species for warming. This idea of a key set of responsive species was partially supported by our data, since both sites had community changes along PCoA1, which could be explained by temperature. However, there were also temperature changes along PCoA2, that were specific for the grassland soil, and thus the overlap in indicator species was relatively small. This suggests that a large part of the warming response will be soil-specific, presumably due to the community filtering effects of other soil physico-chemical characteristics, e.g. pH (Lauber et al., 2009). More comparisons of response patterns across a broader range of soil types are therefore needed. Nevertheless, the use of growth based trait or function determination, as used here for temperature, may be an efficient tool to disentangle driving factors for the community composition (Hicks et al., 2021), as done earlier for pH (Fernández-Calviño and Bååth, 2010), soil salinity (Rath et al., 2019), heavy metals (Fernández-Calviño et al., 2011), or moisture (de Nijs et al., 2019).

Although this geothermic gradient is a natural event and therefore worth studying in its own right, much of the interest in similar areas is related to the need to better understand temperature effects on soil processes in order to predict effects of future global change. It is important to bear in mind that geothermic warming (and similar effects due to heating cables) is not a perfect simulation of a warming climate, as the warming effects are limited to the soil and, to a lesser extent, the immediately overlying vegetation (Sigurdsson et al., 2016). However, for the present study with focus on direct temperature effects on the soil microbes, this drawback is of minor importance, since the soil organisms will be directly affected by temperature irrespective of whether the above ground environment is warmed to the same extent. Thus, our result on bacterial adaptation to warming, the extent of adaptation and the possibility of threshold temperatures for effects will be of interest for prediction and understanding of future global change effects.

## Acknowledgements

Research at the FORHOT site is supported project grant from the Icelandic Research Council (Rannsóknasjóður, ForHot-Forest, Project No 163272-051). Fieldwork for the present study was supported by EU-COST action ClimMani (ES1308) to JTW. We thank the FORHOT research community for constructive and stimulating discussions of the data presented here, and the Lorentz Center, Leiden, NL, for a workshop grant in 2014.. This work was part of LUCCI (Lund University Centre for studies of Carbon Cycle and Climate Interactions).

## Appendix 1 Interpolation and uncertainty propagation of soil temperature data

The plots established along the soil temperature gradients in both the grassland and forest sites were based on snapshot soil temperature measurements using a hand-held probe. In subsequent years, after the samples for the current study were taken, permanent soil temperature sensors and loggers were installed to allow more precise and dynamic soil temperature data. Due to the imprecision of the hand-held temperature measurements, and to allow for the possibility of discrepancy between temperature elevation measured on a given day, and longer-term trends in temperature elevation, it was decided to use the temperature logger data to calculate better estimates of average temperature elevation.

**Figure S 1.**
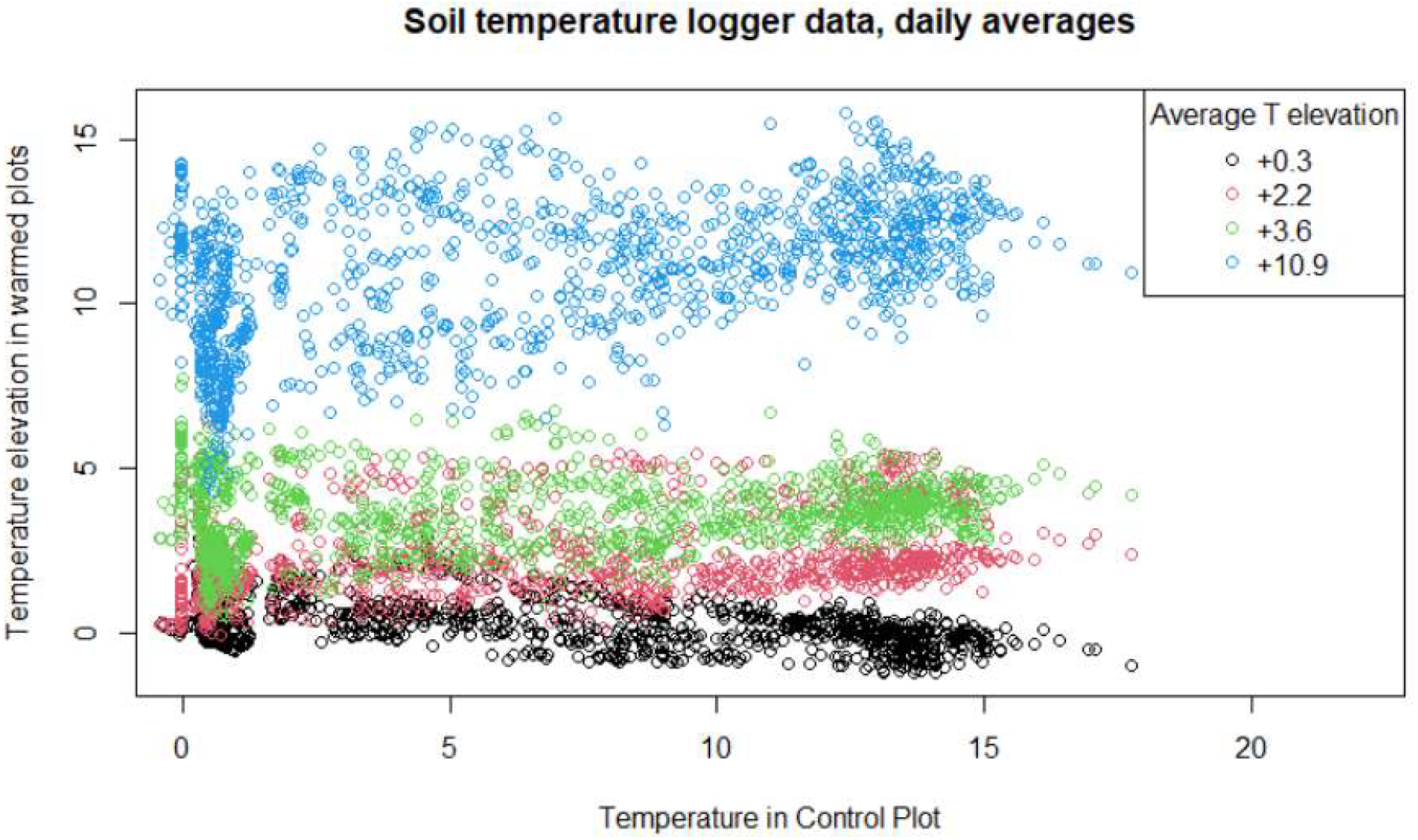
Relationship between temperature elevation and soil temperature in control (ambient) plots over three years of sampling. Each point represents a daily average. The overall tendency of elevation of each plot is relatively consistent throughout the year (no linear trend) but there is considerable day to day variation. This variation was estimated and incorporated into the estimation procedure relating hand-held soil temperature measurements to long-term average elevations.

Logger data from 5 locations in each site were used to model the relationship between daily temperature averages in control plots and for each of the 4 other plots (spanning temperature elevations between +0.3 and +11.0 °C relative to ambient, Figure S1). The estimates of variance as a function of temperature elevation were then combined with the hand-held soil temperature measures (*n* = 1 – 3 different measurements depending on plot) to calculate maximum likelihood estimates and associated errors of the long-term average temperature elevation for each site (Figure S2)

**Figure S 2.**
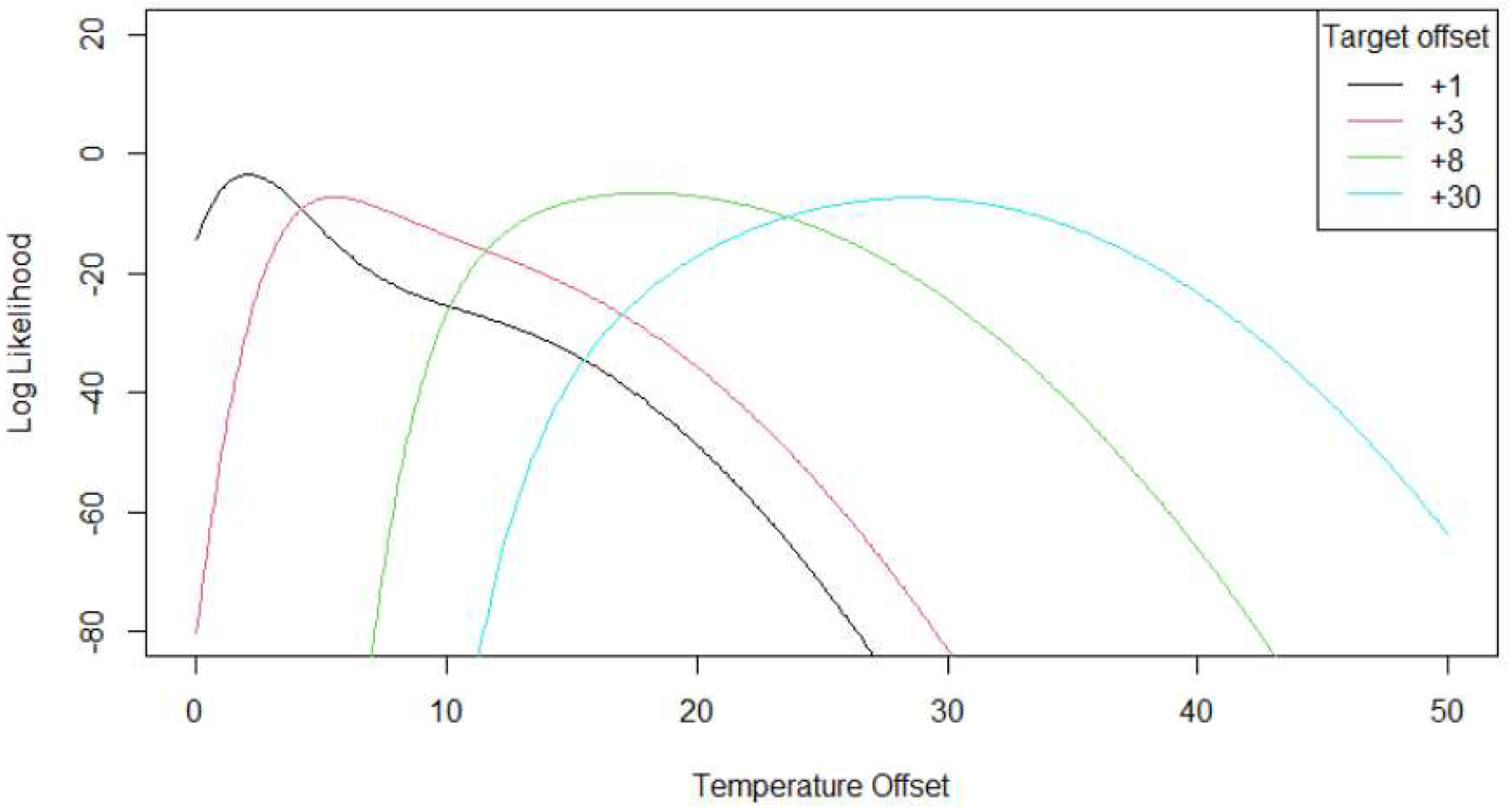
Example of maximum likelihood estimate of temperature offset based on patchy hand-held temperature data. Curves are likelihood profiles for a parameter representing the real (unmeasured) long-term temperature offset. Likelihood profiles were generated by assuming normal errors with variance estimated from the logger data (see Figure S1). Wider profiles indicate greater uncertainty in the estimate of the underlying temperature offset.

## Notes

### Competing Interest Statement

The authors have declared no competing interest.

## References

Anderson, M.J., 2001. A new method for non-parametric multivariate analysis of variance. Austral Ecology 26, 32–46.

Bååth, E., 2018. Temperature sensitivity of soil microbial activity modeled by the square root equation as a unifying model to differentiate between direct temperature effects and microbial community adaptation. Global Change Biology 24. doi:10.1111/gcb.14285

Bååth, E., 1994. Measurement of protein synthesis by soil bacterial assemblages with the leucine incorporation technique. Biology and Fertility of Soils 17. doi:10.1007/BF00337747

Bååth, E., Pettersson, M., Söderberg, K.H., 2001. Adaptation of a rapid and economical microcentrifugation method to measure thymidine and leucine incorporation by soil bacteria. Soil Biology and Biochemistry 33. doi:10.1016/S0038-0717(01)00073-6

Bárcenas-Moreno, G., Brandón, M.G., Rousk, J., Bååth, E., 2009. Adaptation of soil microbial communities to temperature: Comparison of fungi and bacteria in a laboratory experiment. Global Change Biology 15, 2950–2957. doi:10.1111/j.1365-2486.2009.01882.x

Bartram, A.K., Lynch, M.D., Stearns, J.C., Moreno-Hagelsieb, G., Neufeld, J.D., 2011. Generation of Multimillion-Sequence 16S rRNA Gene Libraries from Complex Microbial Communities by Assembling Paired-End Illumina Reads. Appl. Environ. Microbiol. 77, 3846–3852.

Bier, R.L., Bernhardt, E.S., Boot, C.M., Graham, E.B., Hall, E.K., Lennon, J.T., Nemergut, D.R., Osborne, B.B., Ruiz-Gonzalez, C., Schimel, J.P., Waldrop, M.P., Wallenstein, M.D., 2015. Linking microbial community structure and microbial processes: an empirical and conceptual overview. Fems Microbiology Ecology 91.

Birgander, J., Reischke, S., Jones, D.L., Rousk, J., 2013. Temperature adaptation of bacterial growth and 14C-glucose mineralisation in a laboratory study. Soil Biology and Biochemistry 65, 294–303. doi:10.1016/j.soilbio.2013.06.006

Bradford, M.A., Davies, C.A., Wallenstein, M.D., 2008. Thermal adaptation of soil microbial respiration to elevated temperature. Ecology Letters 11, 1316–1327.

Caporaso, J Gregory, Bittinger, K., Bushman, F.D., DeSantis, T.Z., Andersen, G.L., Knight, R., 2010. PyNAST: a flexible tool for aligning sequences to a template alignment. Bioinformatics 26, 266–267. doi:10.1093/bioinformatics/btp636

Caporaso, J. G., Kuczynski, J., Stombaugh, J., Bittinger, K., Bushman, F.D., Costello, E.K., Fierer, N., Pẽa, A.G., Goodrich, J.K., Gordon, J.I., Huttley, G.A., Kelley, S.T., Knights, D., Koenig, J.E., Ley, R.E., Lozupone, C.A., McDonald, D., Muegge, B.D., Pirrung, M., Reeder, J., Sevinsky, J.R., Turnbaugh, P.J., Walters, W.A., Widmann, J., Yatsunenko, T., Zaneveld, J., Knight, R., 2010. QIIME allows analysis of high-throughput community sequencing data. Nature Methods 7, 335–336. doi:10.1038/nmeth.f.303

Cavicchioli, R., Ripple, W.J., Timmis, K.N., Azam, F., Bakken, L.R., Baylis, M., Behrenfeld, M.J., Boetius, A., Boyd, P.W., Classen, A.T., Crowther, T.W., Danovaro, R., Foreman, C.M., Huisman, J., Hutchins, D.A., Jansson, J.K., Karl, D.M., Koskella, B., Mark Welch, D.B., Martiny, J.B.H., Moran, M.A., Orphan, V.J., Reay, D.S., Remais, J. V., Rich, V.I., Singh, B.K., Stein, L.Y., Stewart, F.J., Sullivan, M.B., van Oppen, M.J.H., Weaver, S.C., Webb, E.A., Webster, N.S., 2019. Scientists’ warning to humanity: microorganisms and climate change. Nature Reviews Microbiology. doi:10.1038/s41579-019-0222-5

Corkrey, R., McMeekin, T.A., Bowman, J.P., Ratkowsky, D.A., Olley, J., Ross, T., 2016. The biokinetic spectrum for temperature. PLoS ONE 11. doi:10.1371/journal.pone.0153343

Dacal, M., Bradford, M.A., Plaza, C., Maestre, F.T., García-Palacios, P., 2019. Soil microbial respiration adapts to ambient temperature in global drylands. Nature Ecology and Evolution 3, 232–238. doi:10.1038/s41559-018-0770-5

De Boeck, H.J., Vicca, S., Roy, J., Nijs, I., Milcu, A., Kreyling, J., Jentsch, A., Chabbi, A., Campioli, M., Callaghan, T., Beierkuhnlein, C., Beier, C., 2015. Global Change Experiments: Challenges and Opportunities. BioScience. doi:10.1093/biosci/biv099

De Jonge, C., Radujković, D., Sigurdsson, B.D., Weedon, J.T., Janssens, I., Peterse, F., 2019. Lipid biomarker temperature proxy responds to abrupt shift in the bacterial community composition in geothermally heated soils. Organic Geochemistry 137. doi:10.1016/j.orggeochem.2019.07.006

de Nijs, E.A., Hicks, L.C., Leizeaga, A., Tietema, A., Rousk, J., 2019. Soil microbial moisture dependences and responses to drying–rewetting: The legacy of 18 years drought. Global Change Biology 25. doi:10.1111/gcb.14508

DeAngelis, K.M., Pold, G., Topçuoglu, B.D., van Diepen, L.T.A., Varney, R.M., Blanchard, J.L., Melillo, J., Frey, S.D., 2015. Long-term forest soil warming alters microbial communities in temperate forest soils. Frontiers in Microbiology 6. doi:10.3389/fmicb.2015.00104

Delgado-Baquerizo, M., Oliverio, A.M., Brewer, T.E., Benavent-González, A., Eldridge, D.J., Bardgett, R.D., Maestre, F.T., Singh, B.K., Fierer, N., 2018. A global atlas of the dominant bacteria found in soil. Science. doi:10.1126/science.aap9516

DeSantis, T.Z., Hugenholtz, P., Larsen, N., Rojas, M., Brodie, E.L., Keller, K., Huber, T., Dalevi, D., Hu, P., Andersen, G.L., 2006. Greengenes, a chimera-checked 16S rRNA gene database and workbench compatible with ARB. Applied and Environmental Microbiology 72, 5069–5072. doi:Doi 10.1128/Aem.03006-05

Donhauser, J., Niklaus, P.A., Rousk, J., Larose, C., Frey, B., 2020. Temperatures beyond the community optimum promote the dominance of heat-adapted, fast growing and stress resistant bacteria in alpine soils. Soil Biology and Biochemistry 148. doi:10.1016/j.soilbio.2020.107873

Edgar, R.C., 2013. UPARSE: highly accurate OTU sequences from microbial amplicon reads. Nature Methods 10, 996–.

Edgar, R.C., Haas, B.J., Clemente, J.C., Quince, C., Knight, R., 2011. UCHIME improves sensitivity and speed of chimera detection. Bioinformatics 27, 2194–2200.

Fernández-Calviño, D., Arias-Estévez, M., Díaz-Raviña, M., Bååth, E., 2011. Bacterial pollution induced community tolerance (PICT) to Cu and interactions with pH in long-term polluted vineyard soils. Soil Biology and Biochemistry 43. doi:10.1016/j.soilbio.2011.08.001

Fernández-Calviño, D., Bååth, E., 2010. Growth response of the bacterial community to pH in soils differing in pH. FEMS Microbiology Ecology 73. doi:10.1111/j.1574-6941.2010.00873.x

Hicks, L.C., Frey, B., Kjøller, R., Lukac, M., Moora, M., Weedon, J.T., Rousk, J., 2021. Towards a function-first framework to make soil microbial ecology predictive. Ecology. doi:10.1002/ecy.3594

Ji, M., Kong, W., Jia, H., Delgado-Baquerizo, M., Zhou, T., Liu, X., Ferrari, B.C., Malard, L., Liang, C., Xue, K., Makhalanyane, T.P., Zhu, Y.-G., Wang, Y., Pearce, D.A., Cowan, D., 2022. Polar soils exhibit distinct patterns in microbial diversity and dominant phylotypes. Soil Biology and Biochemistry 166, 108550. doi:https://doi.org/10.1016/j.soilbio.2022.108550

Karhu, K., Auffret, M.D., Dungait, J.A.J., Hopkins, D.W., Prosser, J.I., Singh, B.K., Subke, J.A., Wookey, P.A., Agren, G.I., Sebastia, M.T., Gouriveau, F., Bergkvist, G., Meir, P., Nottingham, A.T., Salinas, N., Hartley, I.P., 2014. Temperature sensitivity of soil respiration rates enhanced by microbial community response. Nature 513, 81–+.

Kirschbaum, M.U.F., 2000. Will changes in soil organic carbon act as a positive or negative feedback on global warming? Biogeochemistry 48. doi:10.1023/A:1006238902976

Kirschbaum, M.U.F., 1995. The temperature dependence of soil organic matter decomposition, and the effect of global warming on soil organic C storage. Soil Biology and Biochemistry 27, 753–760. doi:10.1016/0038-0717(94)00242-S

Lauber, C.L., Hamady, M., Knight, R., Fierer, N., 2009. Pyrosequencing-Based Assessment of Soil pH as a Predictor of Soil Bacterial Community Structure at the Continental Scale. Applied and Environmental Microbiology 75, 5111–5120. doi:Doi 10.1128/Aem.00335-09

Leblans, N.I.W., Sigurdsson, B.D., Vicca, S., Fu, Y., Penuelas, J., Janssens, I.A., 2017. Phenological responses of Icelandic subarctic grasslands to short-term and long-term natural soil warming. Global Change Biology 23. doi:10.1111/gcb.13749

Li, J., Bååth, E., Pei, J., Fang, C., Nie, M., 2021. Temperature adaptation of soil microbial respiration in alpine, boreal and tropical soils: An application of the square root (Ratkowsky) model. Global Change Biology 27. doi:10.1111/gcb.15476

Lin, H., Peddada, S. Das, 2020. Analysis of compositions of microbiomes with bias correction. Nature Communications 11, 1–11. doi:10.1038/s41467-020-17041-7

Lozupone, C., Hamady, M., Knight, R., 2006. UniFrac - An online tool for comparing microbial community diversity in a phylogenetic context. Bmc Bioinformatics 7. doi:Artn 371Doi 10.1186/1471-2105-7-371

Marañón-Jiménez, S., Peñuelas, J., Richter, A., Sigurdsson, B.D., Fuchslueger, L., Leblans, N.I.W., Janssens, I.A., 2019. Coupled carbon and nitrogen losses in response to seven years of chronic warming in subarctic soils. Soil Biology and Biochemistry 134. doi:10.1016/j.soilbio.2019.03.028

Marañón-Jiménez, S., Soong, J.L., Leblans, N.I.W., Sigurdsson, B.D., Peñuelas, J., Richter, A., Asensio, D., Fransen, E., Janssens, I.A., 2018. Geothermally warmed soils reveal persistent increases in the respiratory costs of soil microbes contributing to substantial C losses. Biogeochemistry 138. doi:10.1007/s10533-018-0443-0

Marion, G.M., Henry, G.H.R., Freckman, D.W., Johnstone, J., Jones, G., Jones, M.H., Levesque, E., Molau, U., Molgaard, P., Parsons, A.N., Svoboda, J., Virginia, R.A., 1997. Open-top designs for manipulating field temperature in high-latitude ecosystems. Global Change Biology 3, 20–32.

Martiny, J.B.H., Jones, S.E., Lennon, J.T., Martiny, A.C., 2015. Microbiomes in light of traits: A phylogenetic perspective. Science 350, aac9323–aac9323. doi:10.1126/science.aac9323

Monteux, S., Weedon, J.T., Blume-Werry, G., Gavazov, K., Jassey, V.E.J., Johansson, M., Keuper, F., Olid, C., Dorrepaal, E., 2018. Long-term in situ permafrost thaw effects on bacterial communities and potential aerobic respiration. ISME Journal 12. doi:10.1038/s41396-018-0176-z

Nottingham, A.T., Bååth, E., Reischke, S., Salinas, N., Meir, P., 2019. Adaptation of soil microbial growth to temperature: Using a tropical elevation gradient to predict future changes. Global Change Biology. doi:10.1111/gcb.14502

Nottingham, A.T., Fierer, N., Turner, B.L., Whitaker, J., Ostle, N.J., McNamara, N.P., Bardgett, R.D., Leff, J.W., Salinas, N., Silman, M.R., Kruuk, L.E.B., Meir, P., 2018. Microbes follow Humboldt: temperature drives plant and soil microbial diversity patterns from the Amazon to the Andes. Ecology 99. doi:10.1002/ecy.2482

Nottingham, A.T., Hicks, L.C., Meir, P., Salinas, N., Zimmermann, M., Bååth, E., 2021. Annual to decadal temperature adaptation of the soil bacterial community after translocation across an elevation gradient in the Andes. Soil Biology and Biochemistry 158. doi:10.1016/j.soilbio.2021.108217

O’Gorman, E.J., Benstead, J.P., Cross, W.F., Friberg, N., Hood, J.M., Johnson, P.W., Sigurdsson, B.D., Woodward, G., 2014. Climate change and geothermal ecosystems: Natural laboratories, sentinel systems, and future refugia. Global Change Biology 20. doi:10.1111/gcb.12602

Oliverio, A.M., Bradford, M.A., Fierer, N., 2017. Identifying the microbial taxa that consistently respond to soil warming across time and space. Global Change Biology 23, 2117–2129.

Pettersson, M., Bååth, E., 2013. Importance of Inoculum Properties on the Structure and Growth of Bacterial Communities during Recolonisation of Humus Soil with Different pH. Microbial Ecology 66. doi:10.1007/s00248-013-0208-1

Pettersson, M., Bååth, E., 2003. The rate of change of a soil bacterial community after liming as a function of temperature. Microbial Ecology 46. doi:10.1007/s00248-003-0001-7

Pietikäinen, J., Pettersson, M., Bååth, E., 2005. Comparison of temperature effects on soil respiration and bacterial and fungal growth rates. FEMS Microbiology Ecology 52. doi:10.1016/j.femsec.2004.10.002

Poeplau, C., Kätterer, T., Leblans, N.I.W., Sigurdsson, B.D., 2017. Sensitivity of soil carbon fractions and their specific stabilization mechanisms to extreme soil warming in a subarctic grassland. Global Change Biology 23. doi:10.1111/gcb.13491

Poeplau, C., Sigurdsson, P., Sigurdsson, B.D., 2020. Depletion of soil carbon and aggregation after strong warming of a subarctic Andosol under forest and grassland cover. SOIL 6. doi:10.5194/soil-6-115-2020

Post, E., Alley, R.B., Christensen, T.R., Macias-Fauria, M., Forbes, B.C., Gooseff, M.N., Iler, A., Kerby, J.T., Laidre, K.L., Mann, M.E., Olofsson, J., Stroeve, J.C., Ulmer, F., Virginia, R.A., Wang, M., 2019. The polar regions in a 2°C warmer world. Science Advances 5. doi:10.1126/sciadv.aaw9883

Price, M.N., Dehal, P.S., Arkin, A.P., 2009. FastTree: Computing Large Minimum Evolution Trees with Profiles instead of a Distance Matrix. Molecular Biology and Evolution 26, 1641–1650. doi:10.1093/molbev/msp077

Prosser, J.I., 2012. Ecosystem processes and interactions in a morass of diversity. Fems Microbiology Ecology 81, 507–519. doi:10.1111/j.1574-6941.2012.01435.x

Radujković, D., Verbruggen, E., Sigurdsson, B.D., Leblans, N.I.W., Janssens, I.A., Vicca, S., Weedon, J.T., 2018. Prolonged exposure does not increase soil microbial community compositional response to warming along geothermal gradients. Fems Microbiology Ecology 94, fix174–fix174. doi:10.1093/femsec/fix174

Ranneklev, S.B., Bååth, E., 2001. Temperature-Driven Adaptation of the Bacterial Community in Peat Measured by Using Thymidine and Leucine Incorporation. Applied and Environmental Microbiology 67. doi:10.1128/AEM.67.3.1116-1122.2001

Rath, K.M., Fierer, N., Murphy, D. V., Rousk, J., 2019. Linking bacterial community composition to soil salinity along environmental gradients. ISME Journal 13. doi:10.1038/s41396-018-0313-8

Ratkowsky, D.A., Olley, J., McMeekin, T.A., Ball, A., 1982. Relationship between temperature and growth rate of bacterial cultures. Journal of Bacteriology 149. doi:10.1128/jb.149.1.1-5.1982

Ratkowsky, D.A., Olley, J., Ross, T., 2005. Unifying temperature effects on the growth rate of bacteria and the stability of globular proteins. Journal of Theoretical Biology 233. doi:10.1016/j.jtbi.2004.10.016

Rinnan, R., Michelsen, A., Bååth, E., 2011. Long-term warming of a subarctic heath decreases soil bacterial community growth but has no effects on its temperature adaptation. Applied Soil Ecology 47, 217–220. doi:10.1016/j.apsoil.2010.12.011

Rinnan, R., Rousk, J., Yergeau, E., Kowalchuk, G.A., Baath, E., 2009. Temperature adaptation of soil bacterial communities along an Antarctic climate gradient: predicting responses to climate warming. Global Change Biology 15, 2615–2625. doi:10.1111/j.1365-2486.2009.01959.x

Rousk, J., Frey, S.D., Bååth, E., 2012. Temperature adaptation of bacterial communities in experimentally warmed forest soils. Global Change Biology 18, 3252–3258. doi:10.1111/j.1365-2486.2012.02764.x

Rustad, L.E., Campbell, J.L., Marion, G.M., Norby, R.J., Mitchell, M.J., Hartley, A.E., Cornelissen, J.H.C., Gurevitch, J., Gcte-News, 2001. A meta-analysis of the response of soil respiration, net nitrogen mineralization, and aboveground plant growth to experimental ecosystem warming. Oecologia 126, 543–562.

Sigurdsson, B.D., Leblans, N.I.W., Dauwe, S., Gudmundsdóttir, E., Gundersen, P., Gunnarsdóttir, G.E., Holmstrup, M., Ilieva-Makulec, K., Kätterer, T., Marteinsdóttir, B., Maljanen, M., Oddsdóttir, E.S., Ostonen, I., Peñuelas, J., Poeplau, C., Richter, A., Sigurdsson, P., Van Bodegom, P., Wallander, H., Weedon, J., Janssens, I., 2016. Geothermal ecosystems as natural climate change experiments: The ForHot research site in Iceland as a case study. Icelandic Agricultural Sciences 29, 53–71. doi:10.16886/IAS.2016.05

Todd-Brown, K.E.O., Hopkins, F.M., Kivlin, S.N., Talbot, J.M., Allison, S.D., 2012. A framework for representing microbial decomposition in coupled climate models. Biogeochemistry 109, 19–33.

Verbrigghe, N., Leblans, N.I.W., Sigurdsson, B.D., Vicca, S., Fang, C., Fuchslueger, L., Soong, J.L., Weedon, J.T., Poeplau, C., Ariza-Carricondo, C., Bahn, M., Guenet, B., Gundersen, P., Gunnarsdóttir, G.E.G., Kätterer, T., Liu, Z., Maljanen, M., Marañón-Jiménez, S., Meeran, K., Oddsdóttir, E.S., Ostonen, I., Peñuelas, J., Richter, A., Sardans, J., Sigurðsson, P., Torn, M.S., Van Bodegom, P.M., Verbruggen, E., Walker, T.W.N., Wallander, H., Janssens, I.A., 2022. Soil carbon loss in warmed subarctic grasslands is rapid and restricted to topsoil. Biogeosciences Discussions 2022, 1–25. doi:10.5194/bg-2021-338

Walker, T.W.N., Janssens, I.A., Weedon, J.T., Sigurðsson, B.D., Richter, A., Peñuelas, J., Leblans, N.I.W., Bahn, M., Bartrons, M., De Jonge, C., Fuchslueger, L., Gargallo-Garriga, A., Gunnarsdóttir, G.E., Marañón-Jiménez, S., Oddsdóttir, E.S., Ostonen, I., Poeplau, C., Prommer, J., Radujković, D., Sardans, J., Sigurősson, P., Soong, J.L., Vicca, S., Wallander, H., Ilieva-Makulec, K., Verbruggen, E., 2020. A systemic overreaction to years versus decades of warming in a subarctic grassland ecosystem. Nature Ecology and Evolution. doi:10.1038/s41559-019-1055-3

Walker, T.W.N., Kaiser, C., Strasser, F., Herbold, C.W., Leblans, N.I.W., Woebken, D., Janssens, I.A., Sigurdsson, B.D., Richter, A., 2018. Microbial temperature sensitivity and biomass change explain soil carbon loss with warming. Nature Climate Change 8, 885–+.

Wang, Q., Garrity, G.M., Tiedje, J.M., Cole, J.R., 2007. Naïve Bayesian Classifier for Rapid Assignment of rRNA Sequences into the New Bacterial Taxonomy. Applied and Environmental Microbiology 73, 5261–5267. doi:10.1128/aem.00062-07

Weedon, J.T., Kowalchuk, G.A., Aerts, R., Freriks, S., Röling, W.F.M., van Bodegom, P.M., 2017. Compositional stability of the bacterial community in a climate-sensitive Sub-Arctic Peatland. Frontiers in Microbiology 8. doi:10.3389/fmicb.2017.00317

Wieder, W.R., Allison, S.D., Davidson, E.A., Georgiou, K., Hararuk, O., He, Y., Hopkins, F., Luo, Y., Smith, M.J., Sulman, B., Todd-Brown, K., Wang, Y.-P., Xia, J., Xu, X., 2015. Explicitly representing soil microbial processes in Earth system models. Global Biogeochemical Cycles 29, 2015GB005188. doi:10.1002/2015GB005188

Wieder, W.R., Bonan, G.B., Allison, S.D., 2013. Global soil carbon projections are improved by modelling microbial processes. Nature Climate Change 3, 909.

Wieder, W.R., Sulman, B.N., Hartman, M.D., Koven, C.D., Bradford, M.A., 2019. Arctic Soil Governs Whether Climate Change Drives Global Losses or Gains in Soil Carbon. Geophysical Research Letters 46. doi:10.1029/2019GL085543

Yergeau, E., Bokhorst, S., Huiskes, A.H.L., Boschker, H.T.S., Aerts, R., Kowalchuk, G.A., 2007. Size and structure of bacterial, fungal and nematode communities along an Antarctic environmental gradient. Fems Microbiology Ecology 59, 436–451. doi:DOI 10.1111/j.1574-6941.2006.00200.x

Yergeau, E., Kowalchuk, G.A., 2008. Responses of Antarctic soil microbial communities and associated functions to temperature and freeze-thaw cycle frequency. Environmental Microbiology 10, 2223–2235. doi:10.1111/j.1462-2920.2008.01644.x

Zhang, J., Ekblad, A., Sigurdsson, B.D., Wallander, H., 2020. The influence of soil warming on organic carbon sequestration of arbuscular mycorrhizal fungi in a sub-arctic grassland. Soil Biology and Biochemistry 147. doi:10.1016/j.soilbio.2020.107826

Zhou, J., Deng, Y., Shen, L., Wen, C., Yan, Q., Ning, D., Qin, Y., Xue, K., Wu, L., He, Z., Voordeckers, J.W., Van Nostrand, J.D., Buzzard, V., Michaletz, S.T., Enquist, B.J., Weiser, M.D., Kaspari, M., Waide, R., Yang, Y., Brown, J.H., 2016. Temperature mediates continental-scale diversity of microbes in forest soils. Nature Communications. doi:10.1038/ncomms12083

